# A Novel MiRNA Detection System Combining TWJ-SDA, Multistep L-TEAM, and CRISPR-Cas3

**DOI:** 10.1101/2025.09.25.678453

**Authors:** Riko Miyazaki, Kazuki Aibara, Haruhi Isse, Fumiya Kashiwai, Masashi Takahashi, Azusa Onodera, Yuji Tsunekawa, Yuko Yamauchi, Kazuto Yoshimi, Tomoji Mashimo, Ken Komiya, Takashi Okada

## Abstract

MicroRNAs (miRNAs) serve critical regulatory roles in gene expression and are valuable biomarkers for early disease detection. However, their inherent low concentration in biological fluids poses significant detection challenges. Although traditional methods like real-time quantitative PCR (RT-qPCR) are highly sensitive, they require thermal cycling, limiting their application in point-of-care testing (POCT). Here, we present an isothermal amplification-based miRNA detection system integrating Three-Way Junction (TWJ) formation, Multistep Low-Temperature Amplification (L-TEAM), and CRISPR-Cas3-mediated signal amplification. The integration of the Multistep L-TEAM with the TWJ method achieves high sensitivity, detecting miRNA at concentrations as low as 10 femtomolar within 50 minutes, and effectively distinguishes single-nucleotide mismatches. When CRISPR-Cas3-mediated reaction was integrated, it still proved effective for confirming the presence of the target, but its quantitative reliability requires further optimization. We developed a predictive model using machine learning to facilitate rational optimization of experimental conditions through contribution analysis and to establish a methodology for designing more favorable sequences. The modular nature of our method permits adaptation to diverse miRNA targets without modifications to the fundamental amplification mechanism.

## 1 Introduction

MicroRNAs (miRNAs) are short non-coding RNAs (∼20-25 nucleotides) critically involved in gene regulation, with altered expression profiles strongly associated with various diseases, making them contribute to diagnostic biomarkers [1, 2]. The effective detection of miRNAs is difficult because of their very low concentrations (fmol/L levels) in biological samples, high sequence similarity among family members, and the nature of degradation [2]. Conventional detection techniques like real time quantitative-PCR (RT-qPCR), despite being the standard due to their robustness and sensitivity, are constrained by their requirement for thermal cycling, limiting their suitability for resource-limited or decentralized environments [3].

To address these limitations, isothermal amplification methods such as strand displacement amplification (SDA) [4], exponential amplification reaction (EXPAR) [5], and loop-mediated isothermal amplification (LAMP) [6] have been explored. However, specificity issues persist, especially in distinguishing closely related miRNAs. The three-way junction (TWJ) approach has shown enhanced specificity by preventing non-specific amplification [7]. CRISPR-Cas-mediated detection methods, such as DETECTR [8], SHERLOCK [9], and CONAN [10], have also been developed to improve detection limit and specificity. However, these methods require thermal changes during the reaction. In contrast, low-temperature amplification (L-TEAM) allows specific detection at physiological temperatures [11].

In this study, we report an integrated, isothermal amplification strategy combining the TWJ method [7] with the modified L-TEAM [12] and the CRISPR-Cas3-mediated signal amplification [10], operating consistently at physiological temperatures. This approach was specifically designed to detect three miRNAs (miR-10b-5p, miR-375, and miR-30b-5p) supposed to be elevated biomarkers in glaucoma patients [13–15].

Our detection system initiates with the formation of a stable TWJ structure between the target miRNA, a single-stranded DNA (ssDNA) template, and a helper sequence. L-TEAM subsequently amplifies the DNA signal, which then activates a CRISPR-Cas3 complex, leading to a novel signal amplification technique measurable via fluorescence. In addition, we developed a predictive model using machine learning to facilitate rational optimization of experimental conditions through contribution analysis and to establish a methodology for designing more favorable sequences.

## 2 Reaction

In the conventional method of SDA [4], two-component nucleic acid, target and template, forms a complex, here termed the Non-Junction (NJ) method. In contrast, three nucleic acids, when designed with complementary regions, can form a TWJ complex. Previous studies by Chen *et al*. have demonstrated that TWJ complexes provide enhanced sequence specificity compared to the conventional two-component nucleic acid complexes (NJ), significantly reducing non-specific amplification and potential false positives [7]. The incorporation of the TWJ structure allows for precise target recognition and enhances specificity crucial for diagnostic applications. SDA enables isothermal amplification of ssDNA using DNA polymerase [16] and a nicking endonuclease [17], eliminating the need for thermal cycling. As illustrated in Fig. 1, TWJ complex formation of the target nucleic acid with the helper and corresponding template triggers elongation by DNA polymerase from the 3’ end of the helper strand. Subsequently, the nicking endonuclease specifically recognizes and introduces a nick into the double-stranded DNA (dsDNA), creating a new 3’ end from which the polymerase continues elongation, displacing the formerly elongated strand. Continuous cycling of these reactions results in amplification of the ssDNA signal [4]. While SDA typically requires temperatures above the melting temperature (*T*_*m*_) of the primer DNA, the L-TEAM developed by Komiya. K. et al. enables amplification reactions at physiological temperatures, below the *T*_*m*_ [11].

**Fig. 1:**
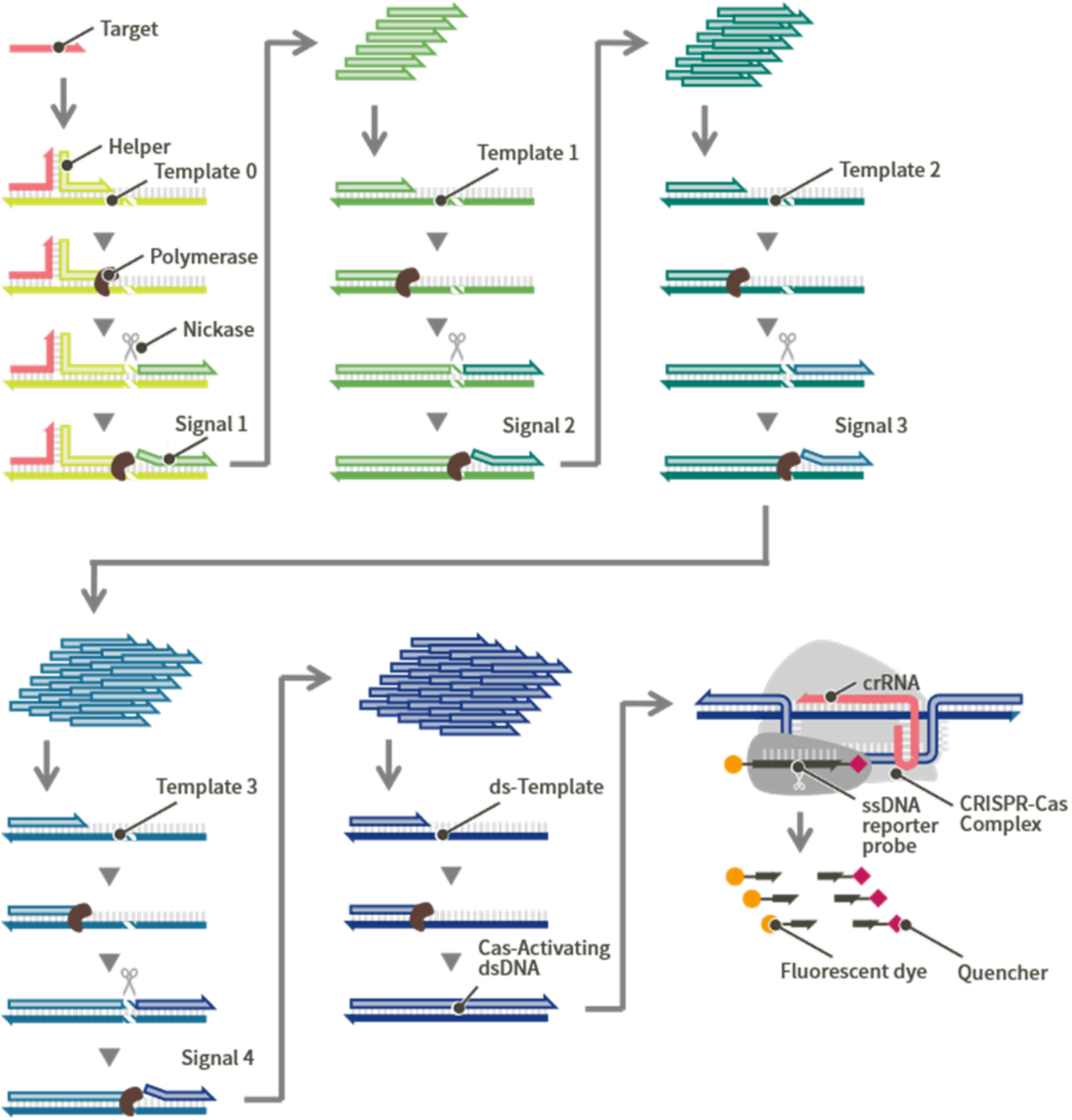
Overview of the mechanism of the proposed detection system. The target hybridizes with the Template 0 and helper ssDNA to form a TWJ complex. The polymerase extends from the 3’ end of the helper, while the nicking endonuclease recognizes and cleaves one strand of the dsDNA. The amplified ssDNA (Signal 1) serves as a primer for the next reaction step, and the successive amplification of Signal 2, 3 and 4 are similarly performed. The Signal 4 hybridizes with the ds-Template, then the polymerase extends from the 3’ end of the signal DNA to produce the Cas-Activating dsDNA. The Cas-Activating dsDNA is not recognized by the endonuclease and instead activates the CRISPR-Cas3 complex, which cleaves the ssDNA region of the ssDNA reporter probe between a fluorescent dye and a quencher.

We further modified and incorporated a sequential amplification method [7], here termed Multistep L-TEAM, to achieve rapid amplification. This approach meets the critical requirements for point-of-care testing (POCT), since it does not require any thermal changes during the reaction [18]. However, the reliance on fluorescence intensity (FI) for evaluating amplification results remains a hurdle for POCT applications. The product from the Multistep L-TEAM process can further activate the CRISPR-Cas3 system, providing additional amplification through the collateral cleavage of a fluorescent reporter [10]. CONAN, developed by Yoshimi et al, integrated LAMP and CRISPR-Cas3-mediated reaction and detected virus RNA by colorimetric change using a lateral flow assay [10, 19]. CrRNA, which functions as a sequence-specific guide molecule, binds to the Cascade complex to recognize dsDNA. The Ecocas3 protein is a bifunctional enzyme that has both 3’ to 5’ ATP-dependent helicase activity and nuclease activity that mainly targets ssDNA. Cas3 and the Cascade/crRNA complex constitute a protein complex (Cas3-Cascade/crRNA). The Cas3-Cascade/crRNA complex activated by dsDNA induces large-scale DNA cleavage extending to the surrounding region of the sequence specified by the crRNA. This property is mainly applied to genome reduction and removal of unnecessary DNA regions [20]. On the other hand, the activated Cas3-Cascade/crRNA complex also has the function of indiscriminately cleaving the surrounding ssDNA through collateral activity. In the present study, this activity was utilized for a different purpose than genetic modification. Metal ion (Mn (II) or Co (II)) is reported to be essential for the CRISPR-Cas3 activity [10]. Specifically, we aimed to induce cleavage of the ssDNA region of the ssDNA reporter probe between the fluorescent dye and quencher dye. This facilitates signal detection and quantification by fluorescence or visual color changes, particularly when integrated into lateral flow assays (LFA) [10, 19].

This innovative combination of Multistep L-TEAM and CRISPR-Cas3 ensures a highly sensitive, specific, and user-friendly system constantly at physiological temperatures, making it ideal for point-of-care diagnostics.

## 3 Materials and Methods

### 3.1 Reagents and Oligonucleotides

Template 1, 2, 3, and ds-Template strands were synthesized by Eurofins Genomics Inc. (Tokyo, Japan) with OPC purification. The molecular beacon (MB) and FQ-Probe were synthesized by Integrated DNA Technologies, Inc. (IDT. Tokyo, Japan) with HPLC purification and modified at its 5’ end with the fluorescent dye 6-FAM and at its 3’ end with the quencher Black Hole Quencher^®^ 1 (BHQ1). SsDNA reporter probe was synthesized by IDT as IDT ZEN^™^ PrimeTime^®^, modified at the 5’ end with the fluorescent dye 6-FAM, at the 3’ end with Iowa Black^®^ Dark Quenchers (IABkFQ), and internally modified with ZEN™ quencher (ZEN) between the ninth and tenth nucleotide from the 5’ end. Other DNA and RNA oligonucleotides used in this study were synthesized by IDT with standard desalting. Bst DNA Polymerase, Large Fragment, Nt.BbvCI nicking endonuclease, Nb.BbvCI nicking endonuclease, 10x rCutSmart buffer, and dNTPs (10 mM) were obtained from New England BioLabs Japan (Tokyo, Japan). SYBR Green II were obtained from Takara Bio Inc. (Shiga, Japan). ATP solution (100 mM) was obtained from Thermo Fisher Scientific (Tokyo, Japan). EcoCas3 protein and the complex of EcoCascade protein and crRNA (Cascade/crRNA) were provided by C4U (Osaka, Japan).

### 3.2 Specificity Evaluation of the TWJ Method

All DNA sequences used in specificity evaluation experiments are listed in Table 1. In addition to the target ssDNA, whose sequence corresponds to the human microRNA, hsa-let-7b and its variants modified at positions 1, 3, 5, 7, 9, 11, 13, and 15 from the 5’ end were synthesized. Reactions were carried out in 20 µL mixtures containing 1x rCutSmart buffer, 10 nM TWJ helper, 10 nM TWJ template, 50 nM hairpin probe, 250 nM FQ-Probe, and either 10 nM target ssDNA or TE buffer as a negative control (NC). As a control experiment using the NJ method, reactions were carried out in 20 µL mixtures containing 1x rCutSmart buffer, 10 nM Duplex template, 50 nM hairpin probe, 250 nM FQ-Probe, and either 10 nM target ssDNA or TE buffer as a negative control (NC). The mixtures also included 0.2 mM dNTP, 0.063 units/µL Bst DNA Polymerase, Large Fragment, and 0.12 units/µL Nt.BbvCI. The reaction mixtures were incubated at 37°C on a real-time PCR system (QuantStudio 3; Thermo Fisher Scientific Inc.), and fluorescence was scanned 600 times at 20-second intervals.

**Table 1:**
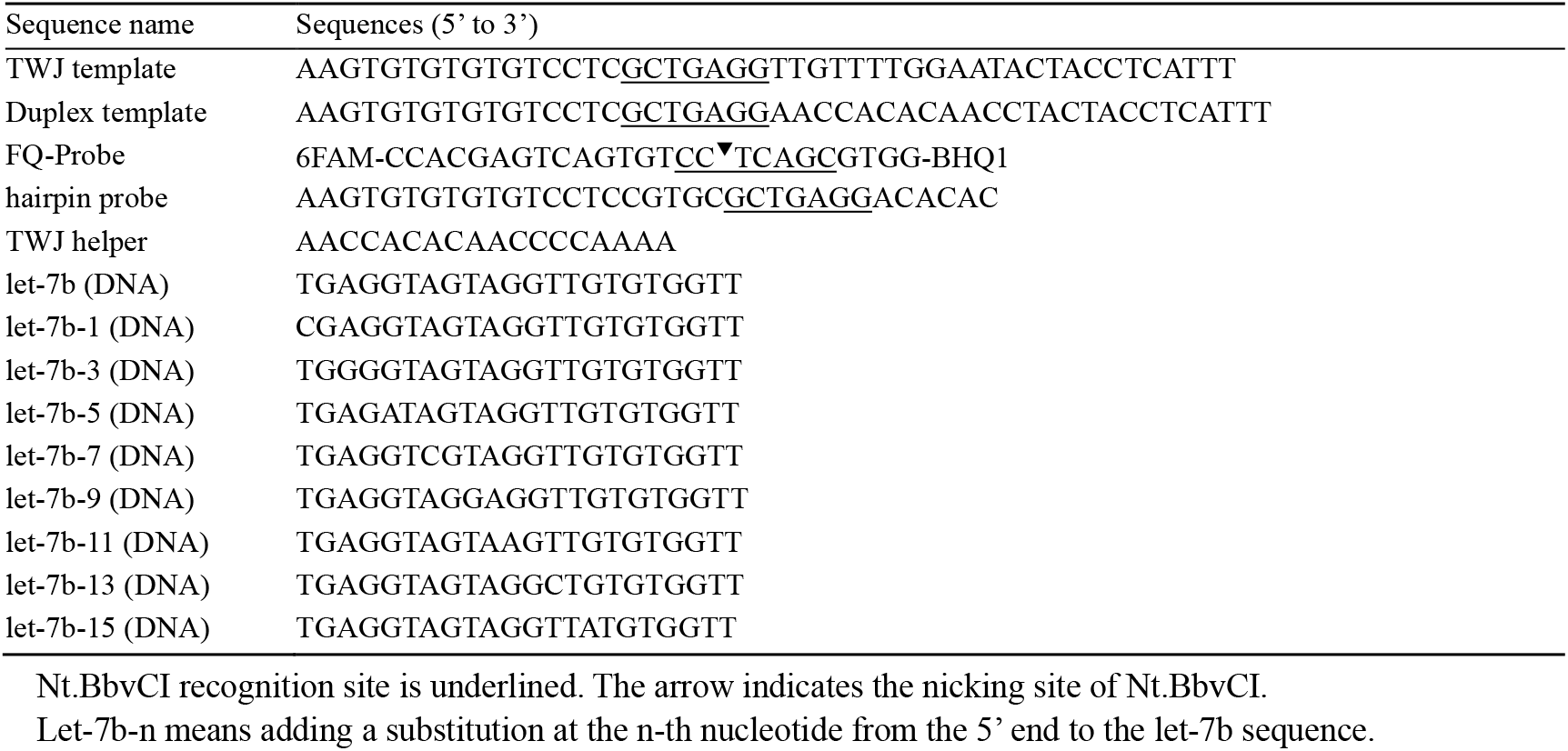
DNA sequences for specificity evaluation

### 3.3 Integration of the TWJ Method with 3-step L-TEAM

Reactions were carried out in 20 µL mixtures containing a custom buffer of our own preparation (60 mM KCl, 10 mM MgCl_2_, 10 nM CoCl_2_, 5 mM HEPES-NaOH [pH 7.7 at 25°C]; since the subsequent experiments were designed as preliminary steps toward the CRISPR-Cas3-mediated reaction, CoCl_2_ required for Cas3 activity was included in the buffer), the 5 nM Template 0, the 10 nM Template 1, the 20 nM Template 2, the 40 nM Template 3, the 5 nM helper, miRNA-10b-5p as target at various concentrations, and the 100 nM MB. The reaction mixture also contained 0.2 mM dNTP, 0.063 units/µL Bst DNA Polymerase, Large Fragment, and 0.12 units/µL Nb.BbvCI. The reaction mixture was incubated at 37°C on the real-time PCR system, and fluorescence was scanned 600 times at 20-second intervals. All sequences used are shown in Table 2.

**Table 2:**
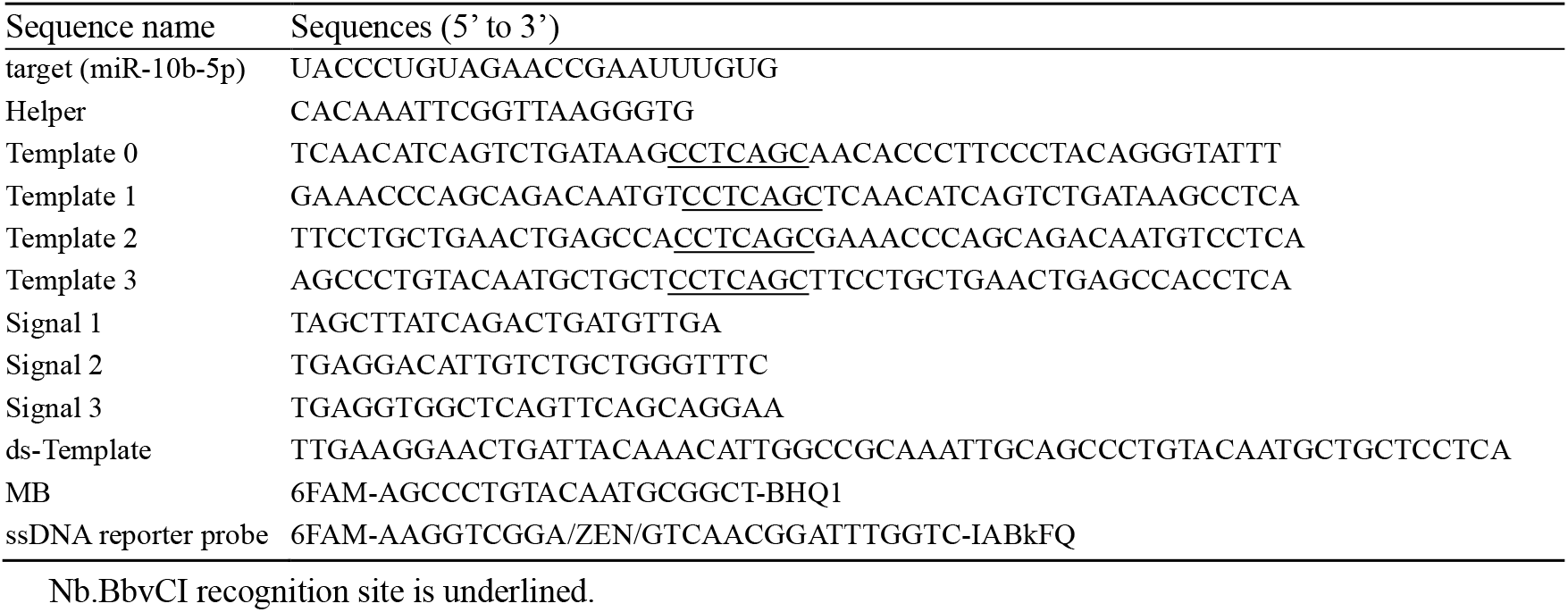
DNA and RNA Sequences of 3-step L-TEAM and CRISPR-Cas3-mediated reaction.

### 3.4 CRISPR-Cas3-Mediated Reactions Following the 3-step L-TEAM

All DNA and RNA sequences used in CRISPR-Cas3-mediated reactions are listed in Table 2. Reactions were carried out in 10 µL mixtures containing the custom buffer, 10 nM Template 1, 20 nM Template 2, 40 nM Template 3, 12 nM ds-Template, Signal 1 as target at varying concentrations, ATP solution (1 mM), and the 500 nM ssDNA reporter probe. The mixtures also included 0.2 mM dNTP, 0.063 units/µL Bst DNA Polymerase, Large Fragment, and 0.12 units/µL Nb.BbvCI. Ecocas3 protein (400 nM), and Cascade/crRNA (100 nM) were further added. The reaction mixture was incubated at 37°C on the real-time PCR system, and fluorescence was scanned 900 times at 10-second intervals.

### 3.5 Sequence Optimization

Target sequences corresponding to the human miRNAs (hsa-miR-10b-5p, miR-375, and miR-30b-5p) were utilized (Table B1 in Appendix B). 12 combinations of the template and helper sequences were designed for each target sequence (designated as targets 1, 2, or 3). The hybridization length between the template and helper was set at 10-12 nucleotides (nt), while that between the template and target miRNA was set at 5-8 nt. Given these parameters and the fixed length of the miRNA, the hybridization length between the helper and miRNA was optimized in fluorescence measurement. FI was measured, and the ratio of FI from reactions with and without the target miRNA as PN ratio was calculated. All sequences used in sequence optimization are provided in Table B1. Reactions were carried out in 20 µL mixtures containing the custom buffer, the 10 nM templates and 10 nM helper, the target miRNA at 10 nM or TE buffer as NC, and SYBR Green II. The reaction mixture also contained 0.2 mM dNTP, 0.063 units/µL Bst DNA Polymerase, Large Fragment, and 0.12 units/µL Nb.BbvCI. The reaction mixture was incubated at 37°C on the real-time PCR system, and fluorescence was scanned 600 times at 20-second intervals.

Machine learning analysis was conducted using Multi-Sigma^®^ software [21], employing binding energies of TWJ structures and reaction times as explanatory variables, with the PN ratio as the objective variable. Optimization, aimed to maximize the PN ratio, are shown in Fig. 2.

**Fig. 2:**
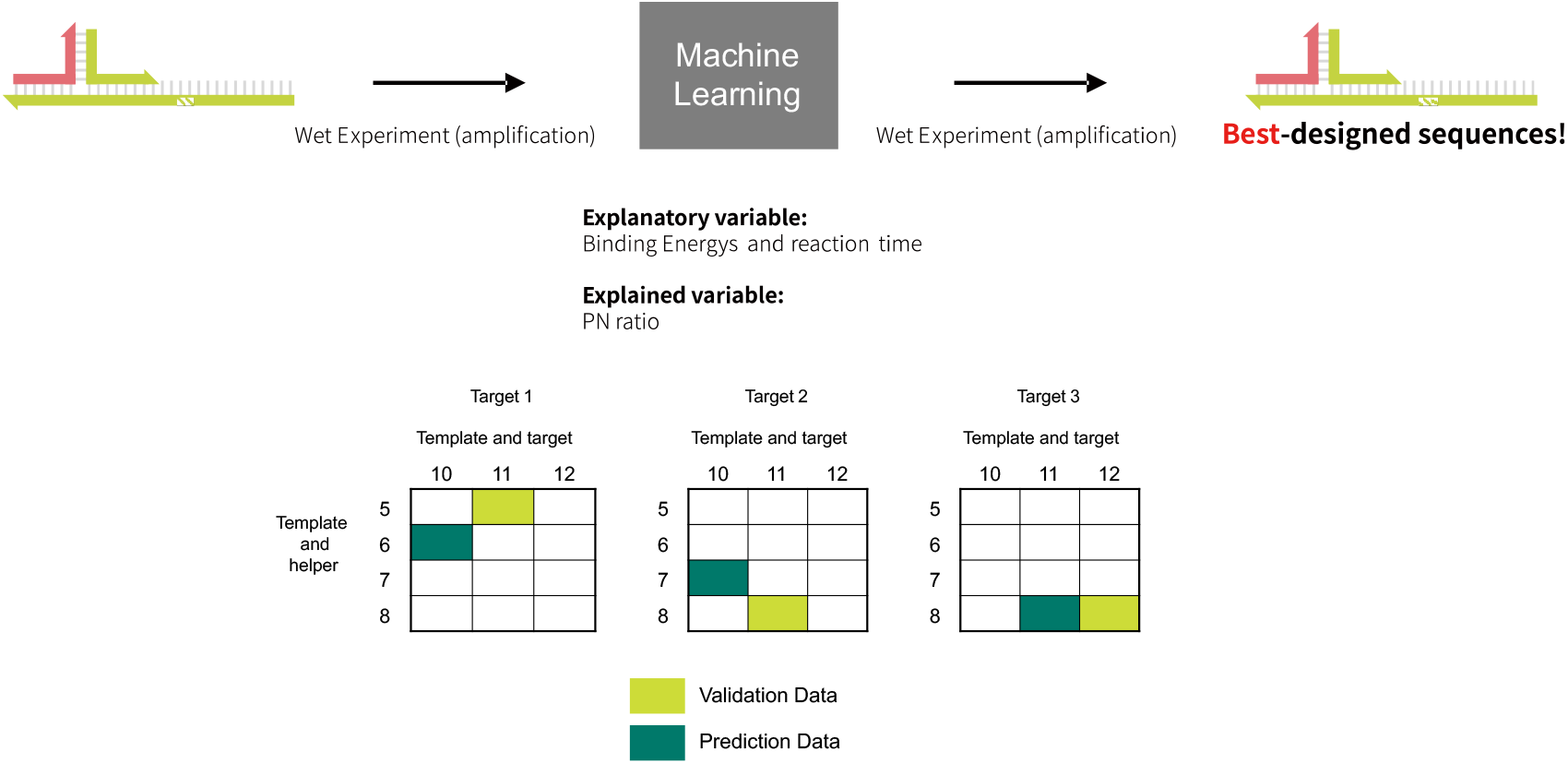
Overview of sequence optimization procedure. Optimization of explanatory variables (binding energies and reaction time) to maximize the experimentally determined PN ratio. Template and helper sequences were subsequently designed based on the optimized binding energies. For each three miRNAs, from 12 combinations of template and helper sequences, one randomly selected combination was used as validation data, and another randomly selected combination was used for prediction.

Nearest-neighbor (NN) model calculations [22, 23] at 37°C were used to estimate binding energies between DNA and RNA. Computer software, NUPACK [24] was used to calculate the binding energies between DNA and DNA. Salt concentration adjustments for binding energies were made according to the methods previously reported by Allawi and SantaLucia [25]. The explanatory variables included four binding energy values and reaction time. Various combinations of four templates and three helper sequences were experimentally tested for each of the three target miRNAs, resulting in a total of 36 sequence combinations. Reaction time was chosen every 20 seconds between 320 and 5980 seconds. The experimentally obtained PN ratio served as the objective variable. The experimental detail was mentioned above.

Experimental data whose PN ratio was below 0.5 were recognized as outliers and not used as learning data. The total number of datasets used as explanatory variables and objective variables was 9501. Using these data, a machine learning model was built based on neural networks via Multi-Sigma^®^ software [21]. For each three miRNAs, from 12 combinations of template and helper sequences, one randomly selected combination was used as validation data, and another randomly selected combination was used for prediction. The remaining ten combinations for each miRNA were used as training data. Consequently, a total of 30 sequence combinations (10 per target) were used for training, 3 sequence combinations were used for validation, 3 sequence combinations were used for prediction. By using the entire dataset belonging to randomly selected sentences for validation and prediction, leakage was supposed to be suppressed as the change of the explained variable over the reaction time was gradual. The PN ratio was predicted from the randomly selected validation sequences and the prediction accuracy was evaluated.

Sensitivity analysis was performed using the Partial Derivative Method (PaD) [26, 27] to estimate the contribution of individual explanatory variables to the objective variable. This method calculates partial derivatives of the output with respect to each input variable, enabling the identification of each explanatory variable’s impact on the predicted outcome. Explanatory variables (binding energies and reaction time) were optimized using a Multi-Objective Genetic Algorithm (MOGA) [28] to maximize the PN ratio (Parameters: population size = 10, generations = 100, crossover rate = 0.6, mutation rate = 0.2, elite rate = 0.2, diversity coefficient = 0.01). Optimal sequences for templates and helper strands were subsequently designed based on neural network predictions using the binding energy calculated by the NN model.

## 4 Result and Discussion

### 4.1 Specificity Evaluation of the TWJ Method

Target specificity of the TWJ method (Fig. 3(A)) and that of the NJ method (Fig. 3(B)) were compared using perfectly matched target sequences and one-nucleotide substituted sequences by measuring FI using the Hairpin probe and FQ-probe (Fig. 3(D)). In the TWJ method, the FI value measured at 40 min for a perfectly matched target sequence was significantly higher (p < 0.05) than that for all substituted target sequences (Fig. 4(A)). Amplification caused by the mismatched target sequences was suppressed by using the TWJ method. In contrast, in the NJ method, the FI value measured for the perfectly matched target sequences was almost the same or lower than that for the mismatched target sequences. Given the small sample size (n=3 for each sample), the results should be interpreted with caution. However, this finding suggests that the TWJ method may have better target specificity than the NJ method.

**Fig. 3:**
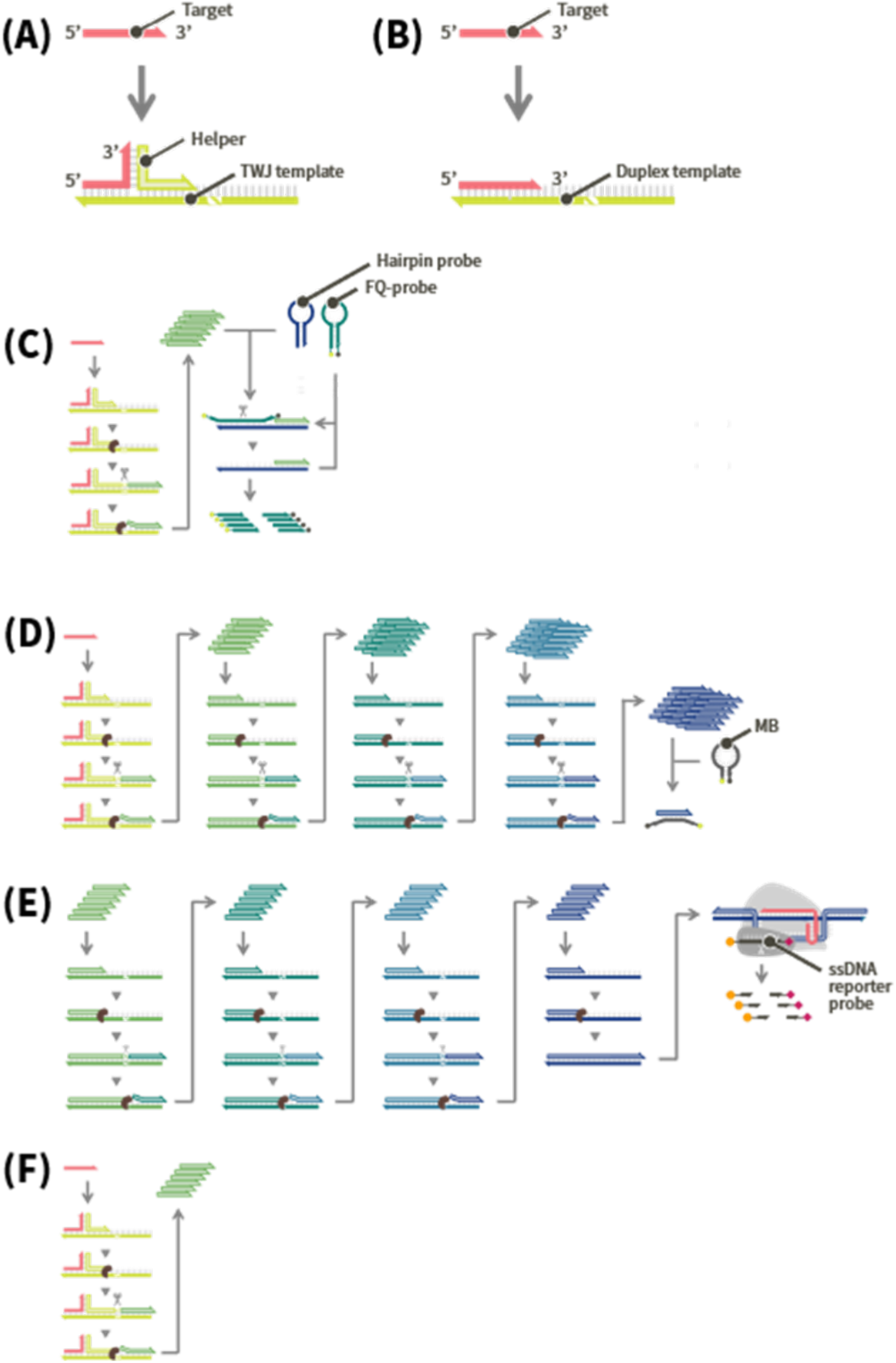
Details of reaction components. Sections 3.2, 3.3, 3.4, and 3.5 provide the experimental conditions for (A)–(F). (A)–(C) correspond to Section 3.2, (D) to Section 3.3, (E) to Section 3.4, and (F) to Section 3.5. (A) The structure of TWJ. The helper and template sequences were designed to form a TWJ structure with the target. (B) The structure of NJ. The target hybridizes with the template. (C) The amplified ssDNA hybridizes with the 5’ end of the hairpin probe (blue), and the 3’ end of the opened hairpin probe hybridizes with the FQ-Probe. The nicking endonuclease then recognizes the dsDNA and cleaves the DNA within the FQ-Probe (emerald green). (D) After the 3-step L-TEAM, the amplified ssDNA hybridizes with the MB and makes its structure change, resulting in fluorescence emission. (E) The final product of the 3-step L-TEAM is converted into dsDNA and integrated with the CRISPR-Cas3-mediated reaction. (F) The amplified ssDNA via the TWJ-SDA was measured using an intercalating dye (SYBR Green II).

**Fig. 4:**
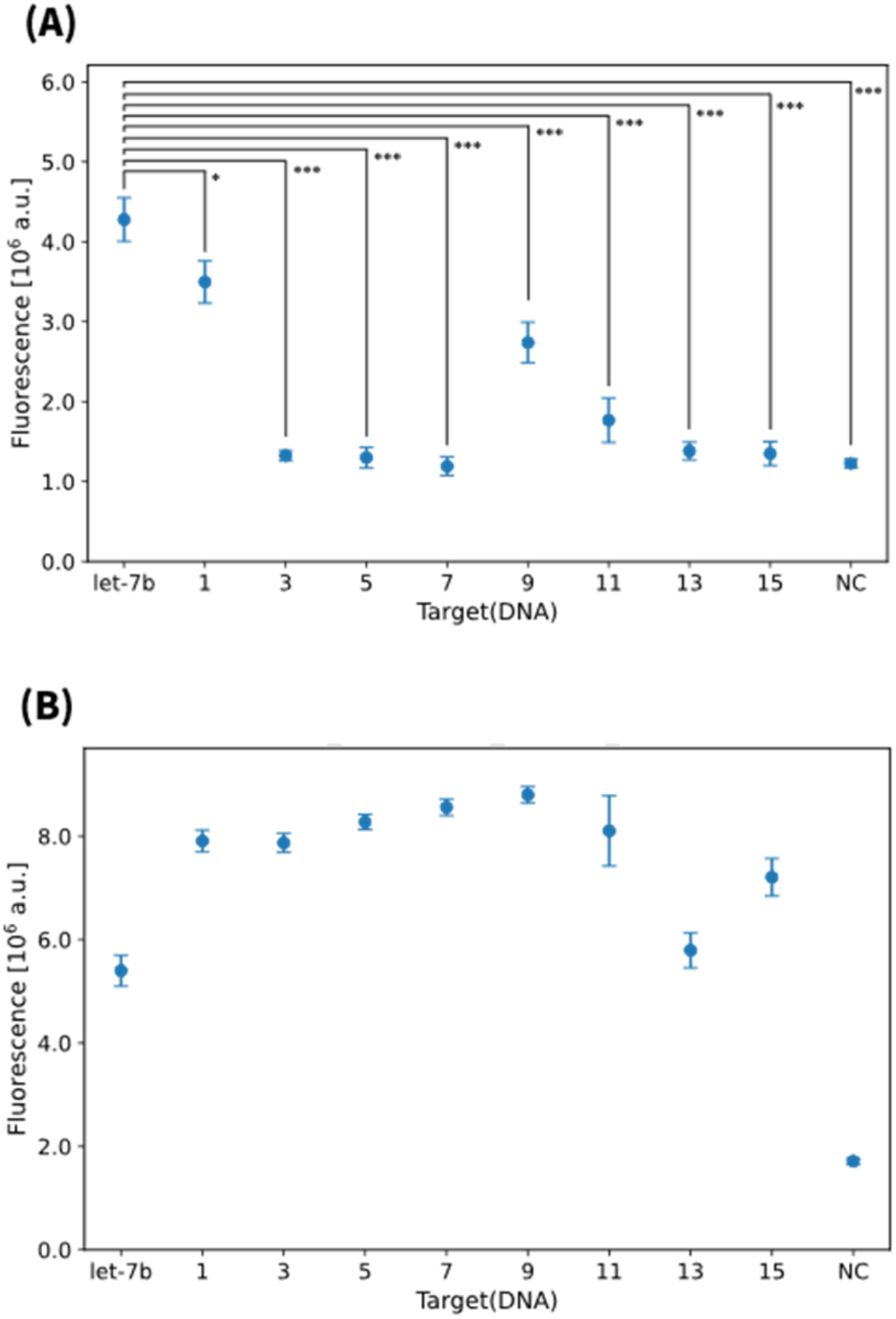
(A) FI at 40 min in the TWJ method. Data are shown as the mean ± s.d. (n=3, FI for the perfect match sequence was significantly higher: *p < 0.05, **p < 0.01, ***p < 0.005, Welch t-test) (B) FI at 40 min in the NJ method. The horizontal axis represents the position of base substitution in the let-7b sequence from the 5’ end.

We further analyzed the FIs to investigate the relationship between the FIs and the thermodynamic properties of the TWJ structure. For the TWJ method, the PN ratio, represented as [the FI measured for each target sequence] / [the FI measured for NC], was calculated and used as a parameter to indicate the amplification efficiency. The thermodynamic properties, Gibbs free energy change caused by the TWJ structure formation, Δ*G*, were calculated using NUPACK [24]. Table 3 showed that substituted target sequences with similar values, comparable to those of the perfectly matched sequences, exhibited higher PN ratios. In contrast, the substituted target sequences with the larger Δ*G*, namely, substitutions that lead to unstable formation, showed the PN ratios around 1, almost the same as the FI of NC. This result suggests that the thermodynamic properties of the TWJ formation play an important role in the target-specific amplification.

**Table 3:** The PN ratios and Δ*G* values calculated for the respective base-substituted positions. Δ*G* represents the Gibbs free energy change upon formation of the TWJ structure, calculated using NUPACK [24].

| Substituted position from the 5' end | PN ratio | $\Delta G$ [kJ/mol] |
| --- | --- | --- |
| None (let-7b, DNA) | 3.110 | -31.57 |
| 1 (let-7b-1, DNA) | 2.371 | -31.39 |
| 3 (let-7b-3, DNA) | 1.064 | -26.40 |
| 5 (let-7b-5, DNA) | 1.061 | -26.82 |
| 7 (let-7b-7, DNA) | 0.970 | -28.45 |
| 9 (let-7b-9, DNA) | 1.775 | -29.71 |
| 11 (let-7b-11, DNA) | 1.284 | -30.08 |
| 13 (let-7b-13, DNA) | 1.120 | -27.89 |
| 15 (let-7b-15, DNA) | 1.085 | -27.10 |

### 4.2 Integration of the TWJ Method with 3-step L-TEAM

The detection limit of the reaction integrating the TWJ method with Multistep L-TEAM (Fig. 3(D)) was assessed with miR-10b-5p as the target at various concentrations under the optimized conditions, including the appropriate design of Template 0, helper strands, and concentration of the polymerase and the nicking endonuclease. A reaction time of 50 min was determined to be optimal, as it resulted in the highest PN ratio for discriminating between the presence and absence of 100 fM target (Fig. C1 and Table C1). Fig. 5 shows the correlation between FI and miR-10b-5p concentrations. A linear relationship exists between FI and the logarithm of miR-10b-5p concentrations, ranging from 10 fM to 1 pM was observed. A previous study on TWJ method [7] demonstrated the linear correlation between FI and miRNA concentrations within 1-50 pM. In this study, the integration of the Multistep L-TEAM with the TWJ method enhanced sensitivity, allowing quantitative detection at substantially lower miRNA concentrations.

**Fig. 5:**
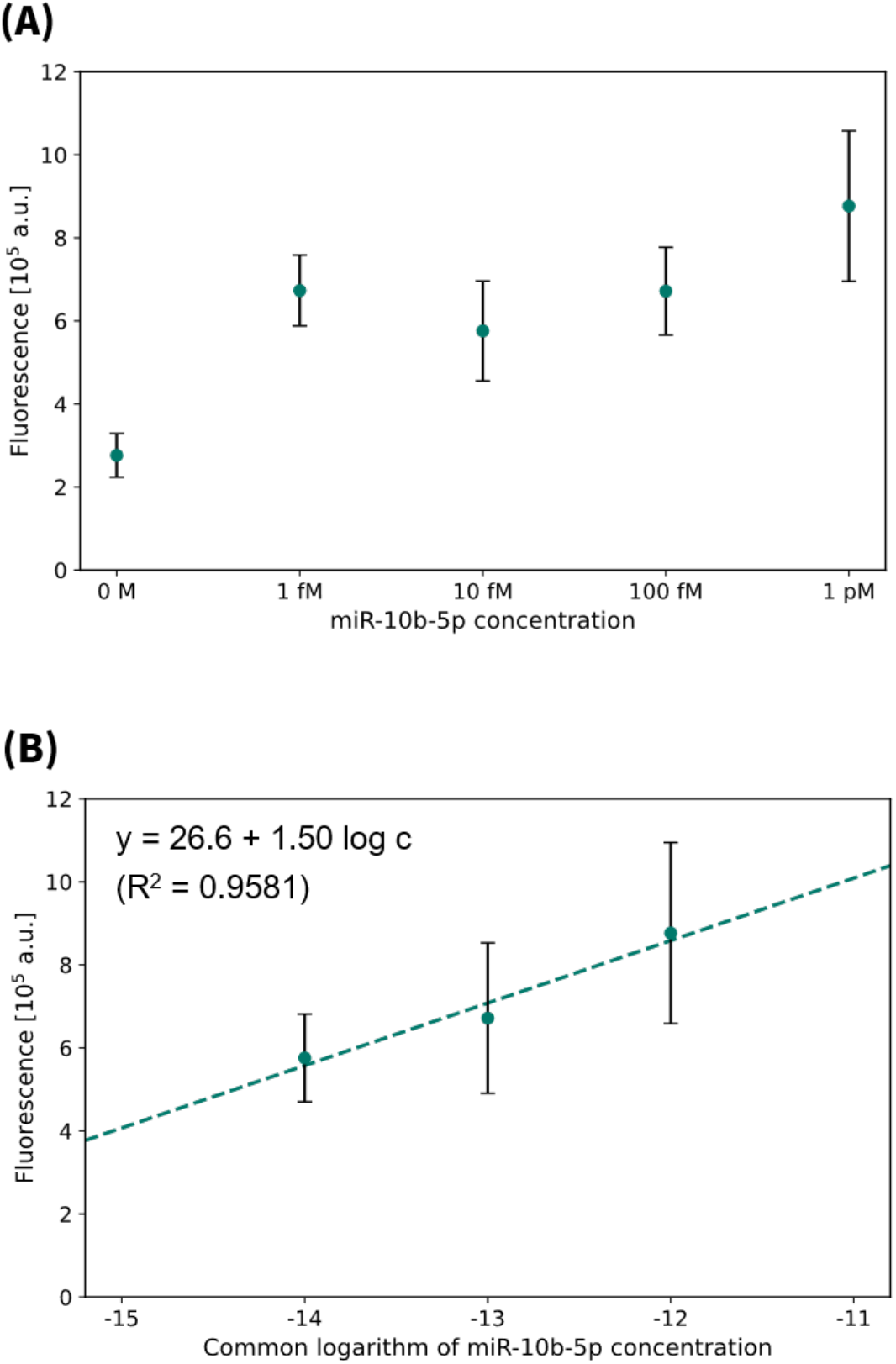
(A) FI at 50 min in the reaction integrating the TWJ method with 3-step L-TEAM. Data are shown as the mean ± s.d. (n=3) (B) linear regression between the common logarithm of miR-10b-5p concentration and FI obtained after 50 min. Data are shown as the mean ± s.d. (n=3)

### 4.3 CRISPR-Cas3-mediated Reaction Following the 3-step L-TEAM

We further integrated the 3-step L-TEAM with the CRISPR-Cas3-mediated reaction (Fig. 1). The detection performance was investigated using 10 nM Template 1, 20 nM Template 2, 40 nM Template 3, and 12 nM ds-Template. Amplification profiles obtained in a single experiment are shown in Fig. 6. The presence or absence of the target could be distinguished, but differences in concentration within the range of 1 pM to 1 fM could not be quantitatively identified. Although the reaction integrating the TWJ method with the 3-step L-TEAM, as described in Section 4.2, achieved quantification of target concentrations, the reaction integrating 3-step L-TEAM with the CRISPR-Cas3-mediated reaction led to non-specific amplification and diminished quantification accuracy (Fig. D1). These results indicated that merely integrating the component reactions was insufficient, highlighting the necessity for further optimization of reaction conditions.

**Fig. 6:**
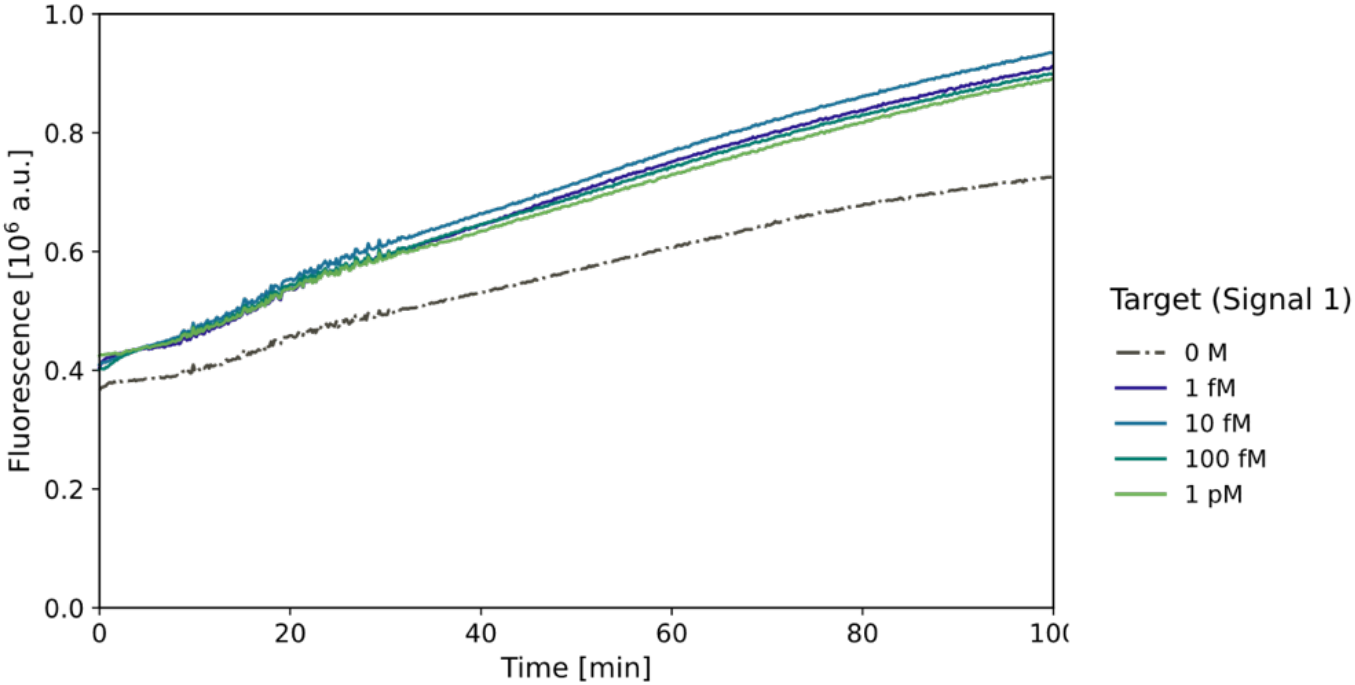
Amplification profiles in the reaction integrating the 3-step L-TEAM with the CRISPR-Cas3-mediated reaction.

### 4.4 Sequence Optimization

We aimed to find the optimal sequences of the template and helper that consist of TWJ. The template sequences were designed to include 10-12 nt sequences complementary to the 5’ end of the target miRNA, and 5-8 nt sequences complementary to the 3’ end of the helper DNA. The 5’ end of the helper sequences were also designed to include sequences complementary to the 3’ end of the target miRNA. Using these sequences, we measured the FI values by using the intercalator dye (SYBR Green II),and calculated the PN. The PN ratios, as the learning datasets employed for the machine learning analysis are summarized in Table 4. While a stable TWJ structure is expected to enable rapid amplification in the presence of the target, it may also lead to target-independent amplification. The optimal binding strength varied depending on the target miRNA. Since the binding affinity depends not only on the length but also on sequence composition, we conducted contribution analysis using binding energies as parameters.

**Table 4:**
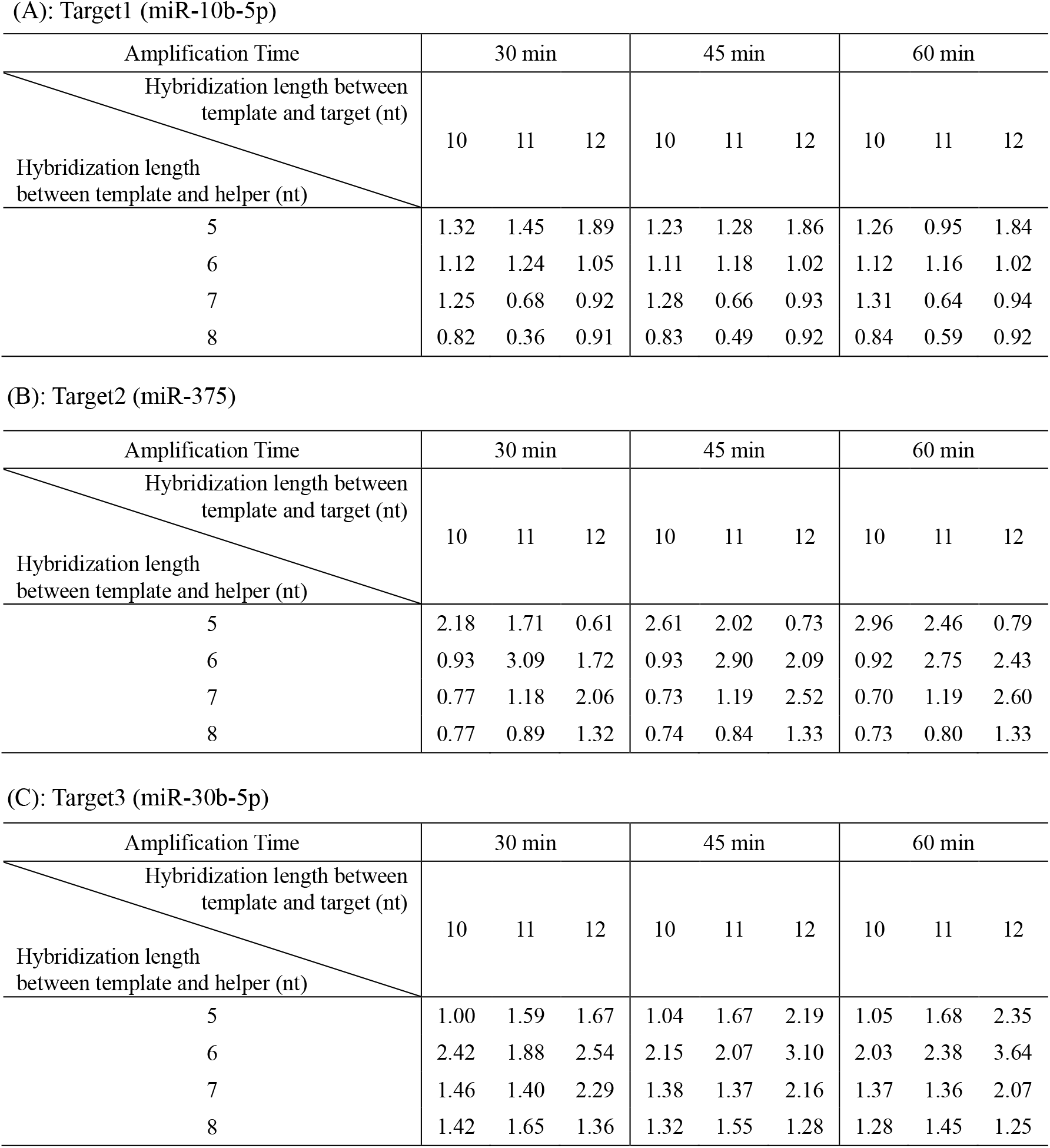
PN ratios after 30, 45, 60 min for various hybridization lengths between template and target, template and helper.

The result of the contribution analysis is shown in Table 5. The binding lengths between the template and the helper were relatively short (5-8 nt), compared to other binding interactions, yet their contribution to specificity was significant. The optimal sequences were designed based on the binding energies. The optimized binding energies, reaction times and DNA sequences are summarized in Table 6. The “optimized” value refers to the binding energy determined by MOGA, while “design” represents the binding energy of sequences designed to closely resemble the optimized value. By utilizing these sequences, we expected to further enhance the PN ratio, achieving highly sensitive amplification while suppressing nonspecific amplification.

**Table 5:**
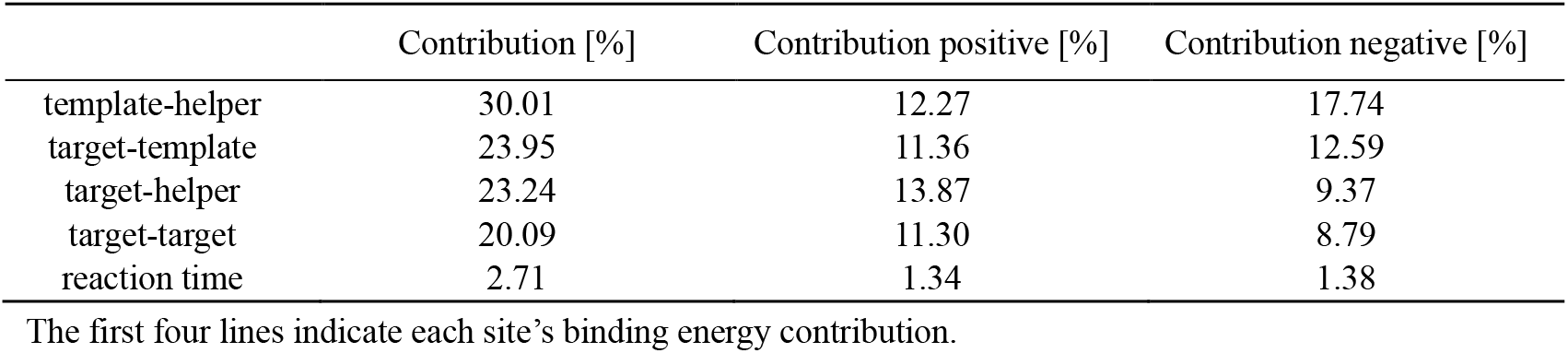
Contribution analysis of PN ratio

|  | Contribution [%] | Contribution positive [%] | Contribution negative [%] |
| --- | --- | --- | --- |
| template-helper | 30.01 | 12.27 | 17.74 |
| target-template | 23.95 | 11.36 | 12.59 |
| target-helper | 23.24 | 13.87 | 9.37 |
| target-target | 20.09 | 11.30 | 8.79 |
| reaction time | 2.71 | 1.34 | 1.38 |

The first four lines indicate each site's binding energy contribution.

**Table 6:** Optimized parameters and designed sequences. Nb.BbvCI recognition site is underlined.

(A): Target1 (miR-10b-5p)
|  | reaction time<br>(sec) | target-template<br>(kJ/mol) | target-helper<br>(kJ/mol) | template-helper<br>(kJ/mol) | target-target<br>(kJ/mol) |
| --- | --- | --- | --- | --- | --- |
| optimized | 4575 | -10.10 | -9.45 | -7.30 | -20.76 |
| design |  | - 9.50 | -8.80 | -7.29 | -20.76 |

|  | reaction time<br>(sec) | target-template<br>(kJ/mol) | target-helper<br>(kJ/mol) | template-helper<br>(kJ/mol) | target-target<br>(kJ/mol) |
| --- | --- | --- | --- | --- | --- |
| optimized | 1642 | -13.99 | -12.39 | -7.45 | -28.55 |
| design |  | -14.00 | -11.70 | -7.48 | -28.55 |

|  | reaction time<br>(sec) | target-template<br>(kJ/mol) | target-helper<br>(kJ/mol) | template-helper<br>(kJ/mol) | target-target<br>(kJ/mol) |
| --- | --- | --- | --- | --- | --- |
| optimized | 5713 | -11.25 | -9.22 | -7.26 | -21.67 |
| design |  | -11.9 | -8.8 | -7.29 | -21.67 |

Whether the PN ratio improved, compared to the data presented in Table 4, was experimentally investigated using the designed sequences. Experimental results, shown in Table 7, indicate that the PN ratios did not significantly surpass those obtained with the original training dataset sequences. Among the three targets tested, miR-10b-5p demonstrated relatively favorable outcomes at the concentration of 10 nM. The results of FI measurement, shown in Fig. 7, exhibited a linear correlation with the logarithm of miR-10b-5p concentrations ranging from 100 fM to 10 pM. However, the PN ratios were not higher than those of a learning dataset. The lack of improvement in the PN ratio highlights the necessity for further optimization. The present data analysis involved the procedures of converting discrete nucleotide sequences into continuous numerical values (binding energies) and identifying theoretically optimal binding energies. Nonetheless, it is inherently difficult to design sequences that exactly match these ideal energy values. Future improvements in optimizing discrete nucleotide sequences may be achieved by integrating surrogate models [29] with binding energy calculation algorithms such as those employed in NUPACK [24].

**Table 7:** Experimental results of the PN ratio for the designed sequence.

| Reaction time | 30 min | 45 min | 60 min |
| --- | --- | --- | --- |
| Target1 (miR-10b-5p) | 1.454 | 1.465 | 1.463 |
| Target2 (miR-375) | 1.211 | 1.218 | 1.219 |
| Target3 (miR-30b-5p) | 1.357 | 1.362 | 1.369 |

**Fig. 7:**
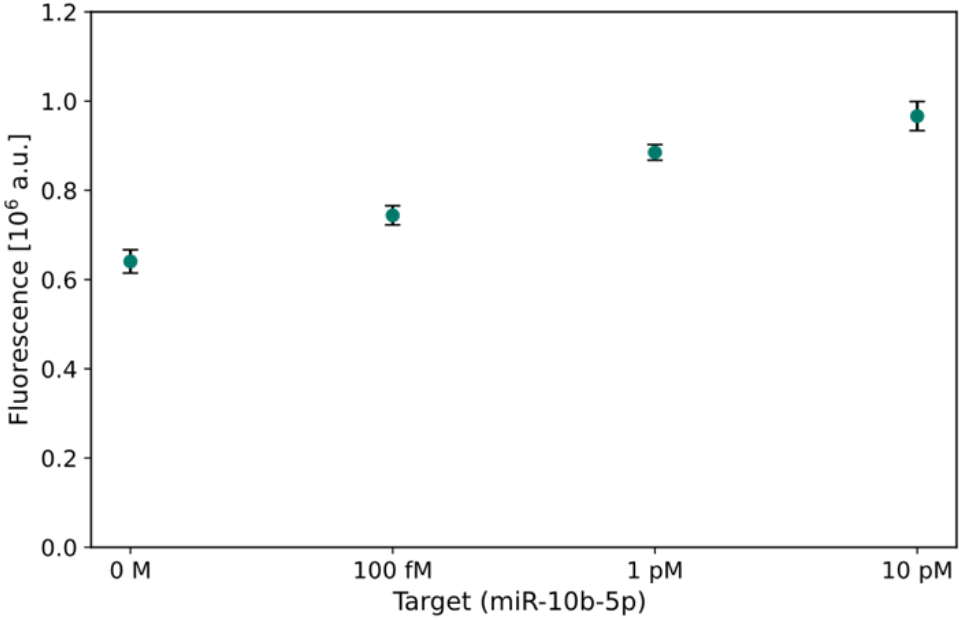
FI after 30 min in the TWJ method experiment with the designed sequences for miR-10b-5p. Data are shown as the mean ± s.d. (n=3).

## 5 Conclusions

In this study, we devised a highly sensitive isothermal miRNA detection system that combines the TWJ method [7], Multistep L-TEAM [12], and CRISPR-Cas3-mediated signal amplification [10]. We obtained proof-of-concept results for combining the L-TEAM and CRISPR-Cas3-mediated reaction, as well as the TWJ method and the Multistep L-TEAM. The present approach facilitated specific and highly sensitive detection of miRNAs as low as 10 fM at a physiological temperature in the reaction combining the TWJ method and the 3-step L-TEAM. The incorporation of the TWJ method significantly improved specificity by reducing nonspecific amplification, while the Multistep L-TEAM efficiently enhanced the overall sensitivity of detection. The presence or absence of the target could be distinguished in the reaction combining the 3-step L-TEAM and CRISPR-Cas3-mediated reaction; however, quantitative amplification could not be achieved. As a consequence, the proposed approach demonstrated modularity and adaptability suitable for detecting different miRNA targets only by designing the template and helper sequences. This modular nature would enable its use in various clinical diagnostic applications. Integration of the present detection system with portable technologies such as lateral flow assays could allow rapid, affordable, and user-friendly POCT [19], potentially transforming early disease detection and monitoring.

## Appendix A. Summary of experimental conditions

**Table A1:** Experimental conditions.

| Experiment | Reaction | Acquisition interval (s) | Cycles | Buffer | Nickase | Fluorescent reagent | Total reaction volume ($\mu\text{L}$) |
| --- | --- | --- | --- | --- | --- | --- | --- |
| 3.2<br>Specificity evaluation of the TWJ method | Fig. 3(A), (B), (C) | 20 | 600 | rCutSmart | Nt.BbvCI | FQ probe | 20 |
| 3.3<br>Integration of the TWJ method with 3-step L-TEAM | Fig. 3(D) | 20 | 600 | The custom buffer | Nb.BbvCI | MB | 20 |
| 3.4<br>CRISPR-Cas3-mediated reactions following the 3-step L-TEAM | Fig. 3(E) | 10 | 900 | The custom buffer | Nb.BbvCI | ssDNA reporter probe | 10 |
| 3.5<br>Sequence optimization | Fig. 3(F) | 20 | 600 | The custom buffer | Nb.BbvCI | SYBR Green II | 20 |
| Appendix D<br>Integration of the final step of L-TEAM and CRISPR-Cas3-mediated reaction | The latter part of Fig. 3(E) | 10 | 900 | The custom buffer | Nb.BbvCI | ssDNA reporter probe | 10 |

## Appendix B. TWJ method

**Table B1:**
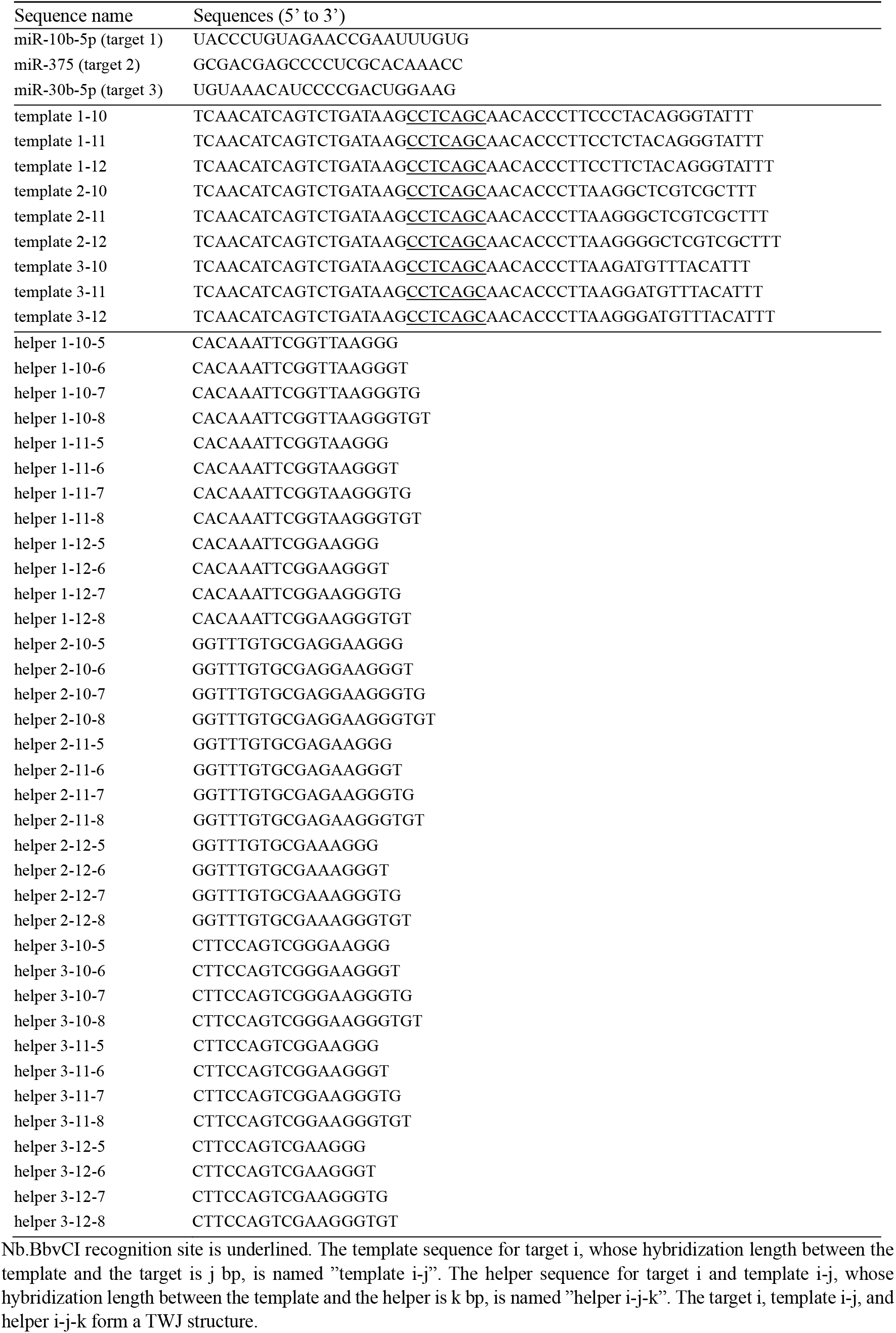
DNA and RNA Sequences for sequence optimization of the TWJ method.

## Appendix C. Integration of the TWJ method with the 3-step L-TEAM

**Fig. C1:**
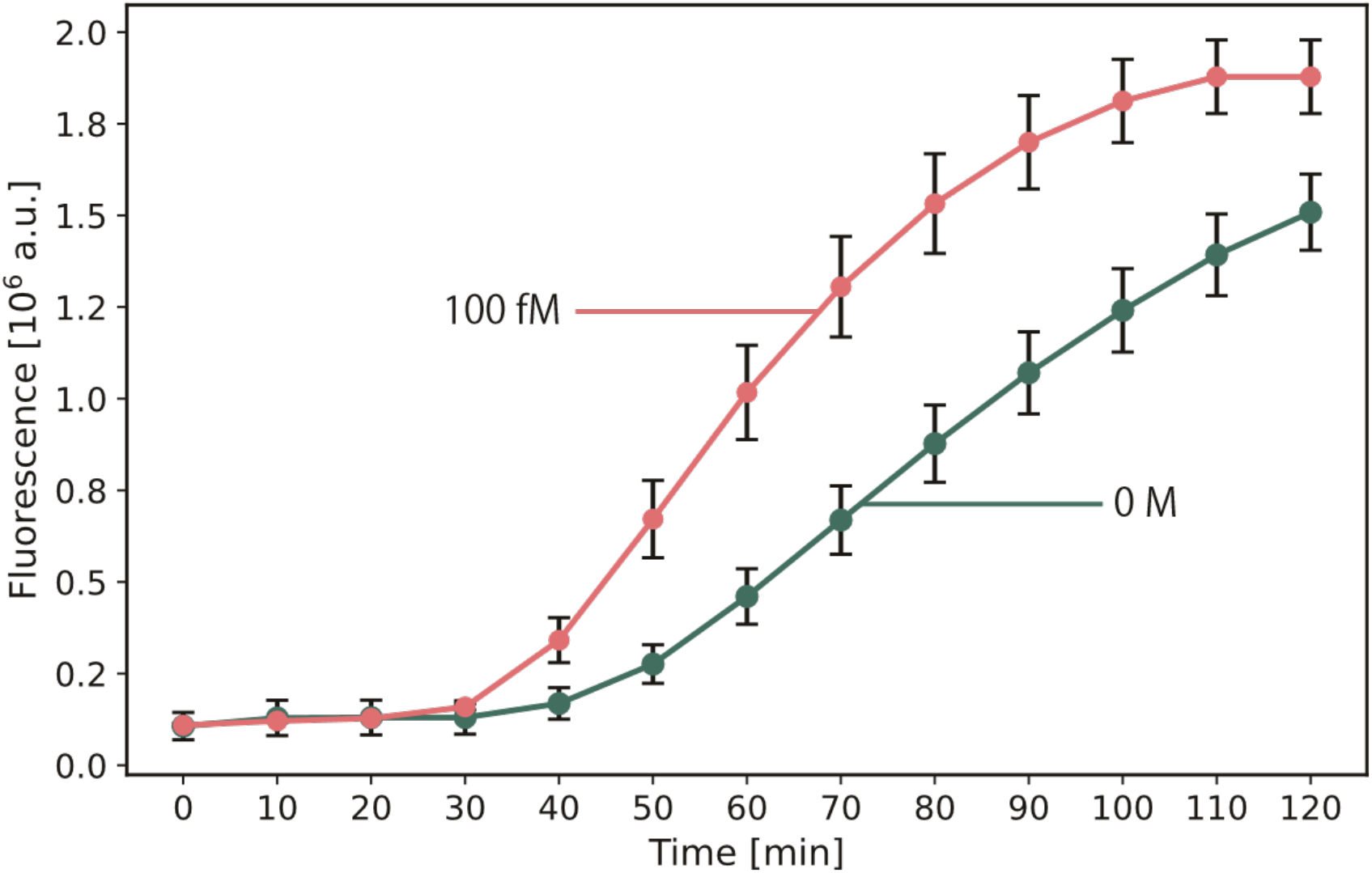
Amplification profiles in the reaction integrating the TWJ method and the 3-step L-TEAM with 100 fM or 0 M miR-10b-5p.

**Table C1:** PN ratios in the reaction integrating the TWJ method and the 3-step L-TEAM.

| Reaction time [min] | 0 | 10 | 20 | 30 | 40 | 50 | 60 | 70 | 80 | 90 | 100 | 110 | 120 |
| --- | --- | --- | --- | --- | --- | --- | --- | --- | --- | --- | --- | --- | --- |
| PN ratio | 1.023 | 0.938 | 0.982 | 1.147 | 2.024 | 2.432 | 2.209 | 1.951 | 1.746 | 1.588 | 1.460 | 1.349 | 1.245 |

PN ratios in the reactions with and without 100 fM are shown. The PN ratio reached the maximum value at 50 min.

## Appendix D. Integration of the final step of L-TEAM and CRISPR-Cas3-mediated Reaction

We integrated the final step of Multistep L-TEAM with CRISPR-Cas3-mediated reaction (Fig. 1). The final product of Multistep L-TEAM, Signal 4, hybridizes with the ds-Template. Elongation of the Signal 4 by DNA polymerase converts the hybridized complex to the Cas-Activating dsDNA, which serves as the activator for the CRISPR-Cas3 complex.

The amplification performance was investigated using the Template 3 with the ds-Template at various concentrations (8, 16, and 24 nM). We prepared 10 µL reaction mixtures in the custom buffer, containing the 40 nM Template 3, the ds-Template, the Signal 3 as target at various concentrations, ATP solution (100 nM) and the 500 nM ssDNA reporter probe. All sequences used in this experiment are listed in Table 2. The reaction mixture also contained 0.2 mM dNTP, 0.063 units/µL Bst DNA Polymerase, Large Fragment, and 0.12 units/µL Nb.BbvCI. Ecocas3 protein (400 nM), and Cascade/crRNA (100 nM) were added last. The reaction mixture was incubated at 37°C on the real-time PCR system, and fluorescence was scanned 900 times at intervals of every 10 seconds.

The amplification profiles with the Signal 3 at the concentrations ranging from 0 to 8 nM are illustrated in Fig. D1. FI were obtained from a single experiment. A trend was observed wherein increased ds-Template concentrations correlated with an accelerated amplification of the NC. Optimal ds-Template concentrations were determined to be either 8 nM or 16 nM, as these concentrations provided a clear distinction between the reaction with the Signal 3 at low concentration and the NC while suppressing non-specific amplification. Assuming that the rate of amplification of the NC is linear with respect to the ds-Template concentrations, we adjusted the ds-Template concentration to 12 nM for subsequent experiments incorporating the 3-step L-TEAM process.

**Fig. D1:**
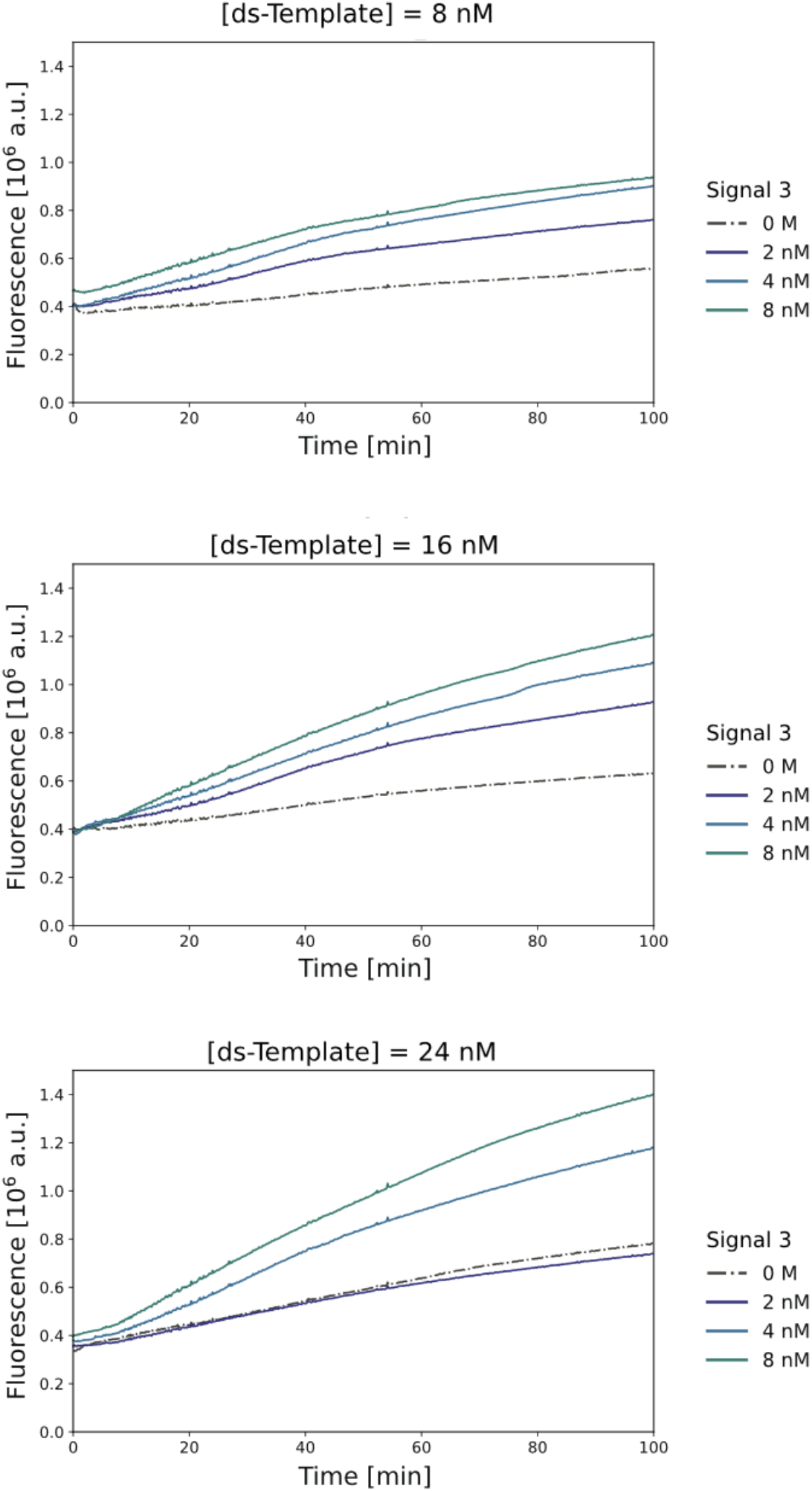
Amplification profiles in the reaction integrating the final step of Multistep L-TEAM with the CRISPR-Cas3-mediated reaction.

## Declarations

The authors received software access and technical advice from AIZOTH Inc.; no author has equity or employment relationship.

## Acknowledgements

This project was conceived as part of a larger project presented at the international Genetically Engineered Machine (iGEM) competition in 2024. Therefore, we would like to thank all the sponsors and the collaborators who supported us. We also thank AIZOTH Inc. for providing financial support and offering their deep learning software, Multi-Sigma^®^. Dr. K. Kawajiri and Dr. R. Kanai of AIZOTH Inc. also kindly provided us with technical help in data analysis using deep learning analysis. Besides, we would like to express our gratitude to everyone who provided personal financial support via the UTokyo Foundation. Finally, we express our sincere thanks to K. Miyamoto, K. Orihara, C. Sakaki, S. Yano, and all members of the iGEM UTokyo.

